# DYRK1A kinase triplication is the major cause of Otitis Media in Down Syndrome

**DOI:** 10.1101/2024.10.03.616443

**Authors:** Hilda Tateossian, Amy Southern, Pratik Vikhe, Eva Lana-Elola, Sheona Watson-Scales, Dorota Gibbins, Debbie Williams, Thomas Purnell, Philomena Mburu, Andrew Parker, Dominic P Norris, Regie Lyn P Santos-Cortez, Brian W Herrmann, Sara Wells, Heena V Lad, Elizabeth MC Fisher, Victor LJ Tybulewicz, Steve DM Brown

## Abstract

Down syndrome (DS), which arises from trisomy of the whole or part of chromosome 21 (Hsa21), is one of the most common genetic abnormalities in humans. DS manifests as a broad spectrum of phenotypic features, including hearing loss due to otitis media with effusion (OME), affecting around 50% of children with DS. We employed a panel of mouse models of DS comprising a nested series of duplications covering the regions of the mouse genome syntenic to Hsa21 in order to define the loci involved with OME in DS. We identified a major locus on mouse chromosome 16, containing only 12 genes, that causes OME. Within this region we demonstrate that normalizing the gene dosage of *Dyrk1a* restored the wild-type phenotype. Investigation of downstream pathways of DYRK1A uncovered a number of pathological mechanisms whereby DYRK1A triplication leads to middle ear inflammation and vascular leak. These include cross-talk of DYRK1A and TGFβ signaling and its impact on proinflammatory cytokines IL-6 and IL-17, as well as raised VEGF levels in the middle ear accompanied by increased *Hif1a*. We conclude that DYRK1A is a potential therapeutic target for OME in children with DS.

## Introduction

Down syndrome (DS) is a common genetic disorder caused by an extra copy of dosage-sensitive genes of human chromosome 21 (Hsa21), and a key challenge remains to determine which genes are causal for aspects of the DS phenotype – the syndrome manifests with a range of physical and intellectual disabilities. More than half of the children with DS have hearing problems as a result of the development of otitis media with effusion (OME)^1^, and this may significantly impact skills such as the acquisition of language. OME is characterised by a middle ear inflammation accompanied by the presence of fluid in the middle ear cavity and thickened middle ear epithelium in the absence of an acute infection. Chronic OME (COME) may last for at least three months. In the general population COME is the most common reason for hearing loss in children and may lead to developmental delay in young children including speech and language impairment. A common treatment for COME is the placement of tympanostomy tubes by surgical procedure into the eardrum ostensibly to prevent the accumulation of fluid in the middle ear and to balance the pressure on each side of the membrane. However, the mechanisms by which tympanostomy tubes reduce inflammation and improve hearing are not clear^2^. For DS, the majority of patients with COME require several rounds of tympanostomy tube placements each involving a surgical procedure with the added risk of complications^3^.

There is a significant genetic component to susceptibility to COME within the human population. Mouse models of COME have elucidated key genes and pathways that may be involved, as well as elaborating the molecular and pathological mechanisms^4–10^. In some cases these loci have been implicated in COME in the human population^11^. However, the genes on human chromosome 21 involved with susceptibility to OME are unknown. In order to identify and characterise the dosage-sensitive genetic loci involved in DS a number of mouse models of DS have been generated and analysed. The mouse orthologues to genes on Hsa21 are spread across three regions of the mouse genome on chromosomes 10 (Mmu10), 16 (Mmu16) and 17 (Mmu17).

An early model, Ts65Dn mice are trisomic for a part of Mmu16^12^ and have been used to study different DS phenotypes, one of which is OME^13^. Auditory-evoked brainstem responses (ABR) were used to assess the hearing of the Ts65Dn mice, demonstrating that the mean ABR thresholds of Ts65Dn were significantly higher than those of the wild-type controls. In addition, histological examination of the ears of mutant mice, aged 3 to 4 months, revealed middle ear inflammation which correlated with the ABR threshold data^13^. However, Ts65Dn animals have a triplication of a large region of Mmu17 containing many genes that do not have synteny to Hsa21, potentially confounding these results^14,15^. Tc1 mice are transchromosomic mice that carry an almost complete copy (75%) of human chromosome 21^16,17^. Auditory function of Tc1 mice was comparable to wild-type animals ^18^. In addition, sections of the middle ears of the mice showed no evidence of fluid in the cavities or thickened middle ear lining. One possible conclusion is that the genes which are trisomic in the Ts65Dn mice (i.e. from the Hsa21-orthologous region of Mmu16) but disomic in Tc1 mice may predispose Ts65Dn mice to OME^18^. A more recent study^19^ examined histological evidence of middle ear inflammation in eight mouse models of DS and found that only Dp(16)1Yey mice, which have full trisomy of the genes from Mmu16, develop significant chronic OME. Similar to the previous study, Tc1 mice did not develop OME and interestingly the authors did not detect signs of middle ear inflammation in Ts65Dn mice. It was concluded that genes trisomic in the Dp(16)1Yey mouse but not in the Tc1 mouse predispose Dp(16)1Yey mice to OM^19^.

Recently, a systematic effort has been underway to create a comprehensive set of genome duplications in the mouse covering those regions of the mouse genome syntenic to Hsa21^20,21^. A set of nested duplications is available that allows for the preliminary localisation of phenotypic effects followed by a more precise refinement of map position to small regions encompassing relatively few genes. This panel of DS mouse models includes Dp1Tyb, Dp(10)2Yey and Dp(17)3Yey encompassing all of the Hsa21- orthologous regions of Mmu16, Mmu10 and Mmu17 respectively. A whole series of smaller nested duplications provide finer resolution across the Dp1Tyb region^20^. These panels of duplications have been employed already to identify regions and loci involved with the heart defects and locomotor dysfunction seen in DS^20,22,23^.

We have employed these panels to undertake a deep genetic analysis of the loci involved with OME in DS. We localised one major locus on Mmu16 that causes the development of highly penetrant OME, and a minor locus, also on Mmu16, which results in low penetrant OME. From within the major locus we found that normalization of gene dosage of the *Dyrk1a* (Dual-specificity tyrosine phosphorylation regulated kinase 1a) gene restored the wild-type phenotype, demonstrating *Dyrk1a* as the causative gene for OME in Down syndrome.

## Results

### Dp1Tyb mouse model of DS displays OME

We examined the presence of otitis media in a number of mouse lines carrying a nested series of genome duplications across chromosome segments syntenic to Hsa21. We first investigated mutants with duplication of the entire syntenic segments from Mmu10 (Dp(10)2Yey), Mmu17 (Dp(17)3Yey) and Mmu16 (Dp1Tyb) (Fig. 1a and Extended Data Fig. 1a). Histological analysis revealed no middle ear inflammation in Dp(10)2Yey or Dp(17)3Yey mice (Extended Data Fig. 1b, Extended Data Table 1). However, all Dp1Tyb mice had characteristic OME: fluid in the middle ear and thickened middle ear lining, from as early as 3 weeks (Fig. 1b). We extended the analysis to mice from different ages (3, 4, 8 and 16 weeks) and from both sexes. All 31 mice we examined had bilateral OME, significantly different from wild-type littermates which could occasionally develop unilateral OME (p < 0.00001) (Extended Data Table 1). Further examination revealed that Dp1Tyb mice also developed a thickened middle ear mucosa. We measured the mucoperiosteal thickness of the middle ears of 3-, 4-, 8- and 16-week old Dp1Tyb and wild-type mice and found significant differences between the two genotypes at all four ages (3wk: p = 0.0010; 4wk: p = 0.0026; 8wk: p <0.0001; 16wk: p = 0.0006) (Fig. 1c).

**Figure 1.**
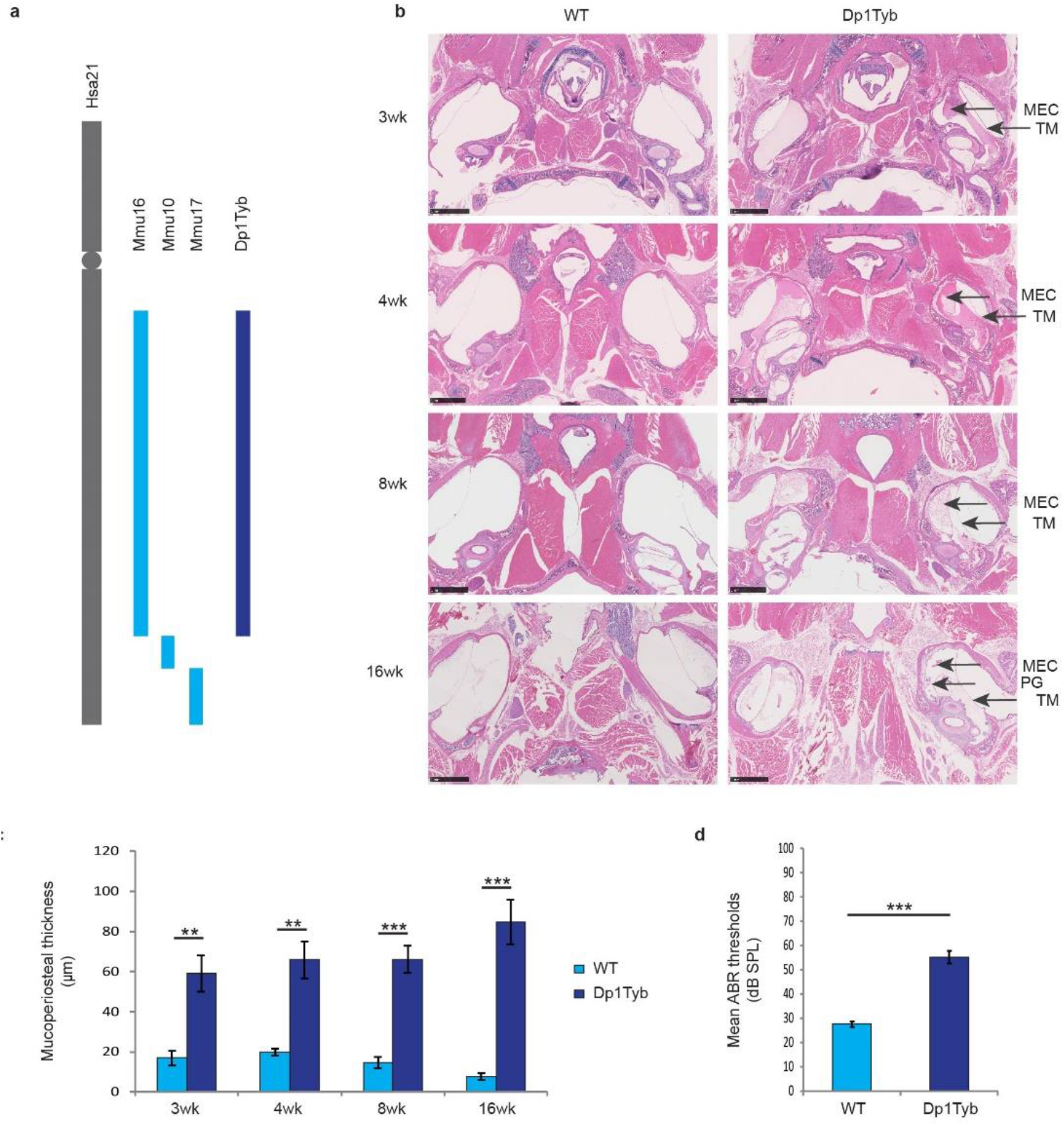
Otitis media phenotype of Dp1Tyb mice. a, Representation of Hsa21 (in grey), the orthologous regions on Mmu10, Mmu17 and Mmu16 (in light blue) and the region of the duplication in Dp1Tyb mice (in dark blue). **b**, Haematoxylin-eosin stained transverse sections through the middle ear of 3 week (3wk), 4 week (4wk), 8 week (8wk) and 16-week-old (16wk) wild-type (WT) and Dp1Tyb mice. MEC: middle ear cavity; TM: tympanic membrane; PG: polypoid growth. Scale bars: 1 mm. **c**, Comparison of the thickness of the epithelial lining of the middle ear for each genotype and age (3wk: WT n = 7 mice (3 female and 4 male), Dp1Tyb n = 7 mice (3 female and 4 male); 4wk: WT n = 5 mice (2 female and 3 male), Dp1Tyb n = 4 mice (2 female and 2 male); 8wk: WT n = 6 mice (3 female and 3 male), Dp1Tyb n = 6 mice (3 female and 3 male); 16wk: WT n = 4 mice (1 female and 3 male), Dp1Tyb n = 7 mice (3 female and 4 male). **d**, Broadband click stimuli ABR thresholds in the right ears of 2-month-old wild-type (WT) and Dp1Tyb mice. The graph shows elevated mean thresholds in Dp1Tyb mice (n = 6 mice, 3 female and 3 male) compared to wild-type (WT) mice (n = 6 mice, 3 female and 3 male). Bars: standard error of mean. P-values were determined using two-tailed t-test. ** p<0.01; *** p<0.001.

We further tested the Dp1Tyb mice for hearing loss by performing click-evoked auditory brainstem response (ABR). At two months of age, the Dp1Tyb carriers had mean thresholds elevated by 27.5 dB SPL compared to wild-type mice (p <0.0001) indicating conductive hearing loss (Fig. 1d). These findings agree with a recent comprehensive phenotyping assessment of Dp1Tyb mice, which also identified raised ABR thresholds^24^. We conclude that there is a locus or loci on mouse chromosome 16 which when duplicated leads to a chronic OME, which is in agreement with observations from previous partial trisomy models of mouse chromosome 16^13^.

### Normal morphology of the cochlea and middle ear in Dp1Tyb mice

The finding of moderately raised ABR thresholds in Dp1Tyb mice accompanied by OME suggests that hearing loss in these mice is caused by a conductive deafness and that sensorineural components in the inner ear are not involved. We sought to confirm this and undertook histological examination of sagittal sections from the cochlea of 4-week old mice, soon after the mice develop OME. Dp1Tyb mice showed no obvious abnormalities of the cochlea (Fig. 2a). In addition, scanning electron microscopy of the organ of Corti at the same age showed normal hair cell and bundle morphology in both mutants and wild-type littermates (Fig. 2b). We conclude that the lack of abnormalities in the inner ear of Dp1Tyb indicates that there is no sensorineural element to the hearing loss in this line. We also assessed if there were abnormalities in the tympanic membrane and ossicles of the middle ear that might contribute to the conductive deafness in Dp1Tyb mice. Visual inspection of the tympanic membrane was used to examine if the membrane was intact. All 8-week old wild-type mice we used for this study (n = 16 ears from 8 mice, 2 female and 6 male) had no visible fluid behind the tympanic membrane, and the malleus (the largest of the three ossicles) was easily recognizable and the membrane was intact. All 8-week old Dp1Tyb mutants analysed (n = 10 ears from 5 mice, 2 female and 3 male) had cloudy ears as a result of the fluid behind the eardrum. The malleus of the mutants was obscured but the membrane was intact (Fig. 2c). μCT analysis of middle ear ossicles in 8-week old mice revealed no difference in the shape and the size of the bones between mutants (n = 8 ears from 4 mice, 2 female and 2 male) and wild-type mice (n = 8 ears from 4 mice, 2 female and 2 male) (Fig. 2d). In summary, we conclude that the hearing loss in Dp1Tyb mice is entirely due to OME within the middle ear.

**Figure 2.**
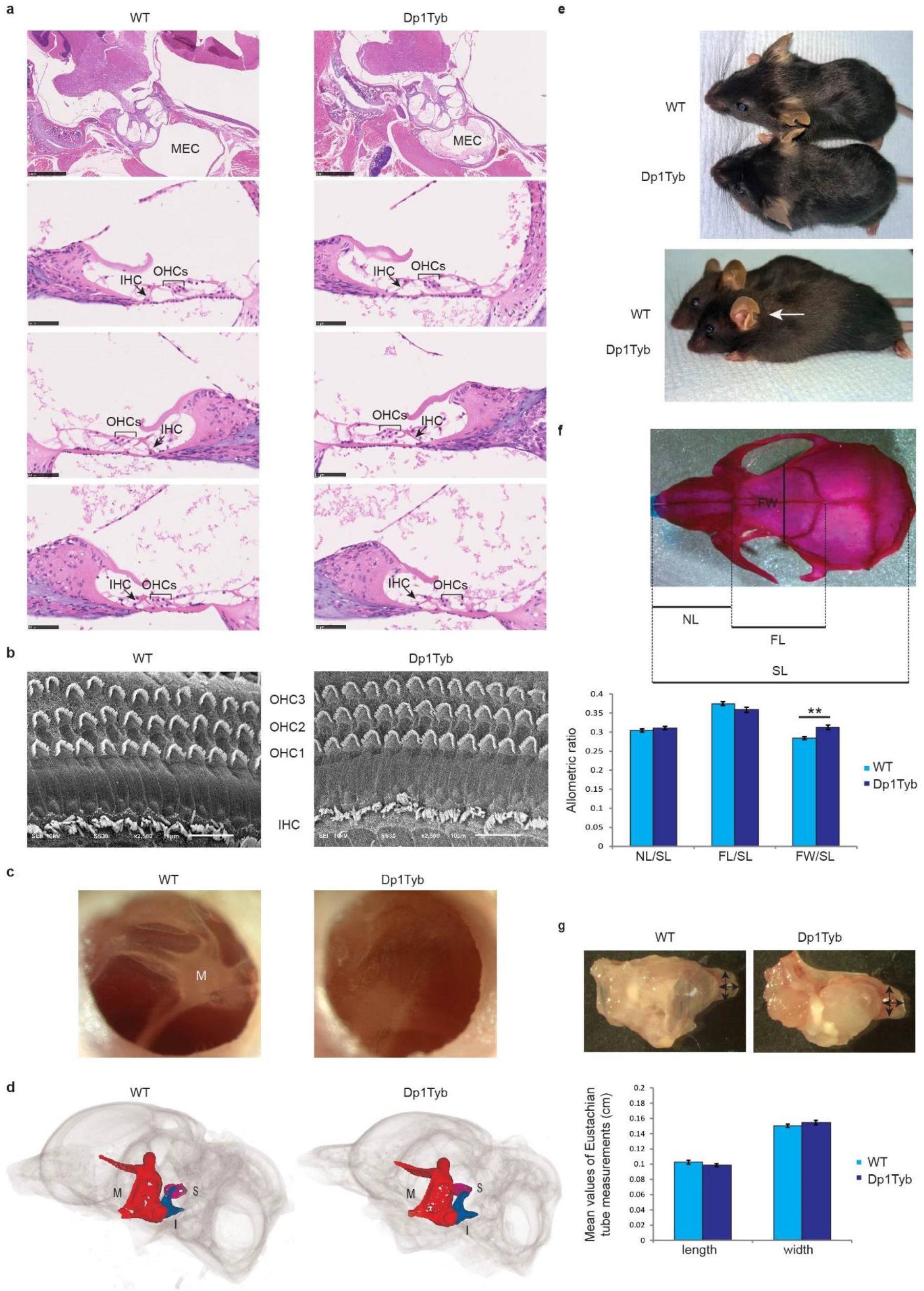
Normal ear morphology and craniofacial dysmorphology in Dp1Tyb mice. **a**, Histological analysis of the hair cells in 2-months old mice from both genotypes. No difference was observed between the genotypes. MEC: middle ear cavity. Scale bars: 1mm and 100 μm. **b**, Scanning electron micrographs showing hair cell organisation and morphology in the mid turn of the cochlea of wild-type (WT) and Dp1Tyb mice. Three rows of OHCs and a row of IHCs were observed in both wild-type and mutant mice. Scale bars: 10 μm. OHC, outer hair cell; IHC, inner hair cell. **c**, Microscopic views of tympanic membrane from wild-type (WT) and Dp1Tyb mice. No visible fluid behind the tympanic membrane and easily recognizable malleus was detected in the wild-type ears and cloudy appearance of the eardrum was detected in all inspected Dp1Tyb middle ears. Both wild-type and mutant ears had intact tympanic membranes. **d**, μCT images of the ear of a wild-type (WT) and a Dp1Tyb mouse showing no visible difference in the size of the malleus (in red), incus (in blue) and stapes (in magenta). M, malleus; I, incus; S, stapes. **e**, Images of a Dp1Tyb mouse and a wild-type littermate showing a different shape of the skull and different shape and position of the pinna in the mutant mouse. The arrow indicates the pinna in the Dp1Tyb mouse. **f**, Dorsoventral view of a 2-month-old wild-type mouse skull showing the measurements used to study the craniofacial defect of the Dp1Tyb mice and graphic comparison of the ratios of the nasal bone length (NL), frontal bone length (FL) and the frontal bone width (FW) to the full dorsal length of the skull (SL) for each genotype. The graph shows significant difference in the width of the Dp1Tyb mice (n = 6 mice, 3 female and 3 male) compared to wild-type mice (WT) (n = 8 mice, 3 female and 5 male). Bars: standard error of mean. P-values were determined using two-tailed t-test ** p<0.01. **g**, Images of dissected ears from wild-type (WT) and Dp1Tyb mice. Arrows indicate the Eustachian tube measurements taken for this study. Comparison of the length and the width of the bony part of the Eustachian tubes in wild- type (WT) and Dp1Tyb mice. Bars: standard error of mean. P-values were determined using two-tailed t-test. No significant difference was detected.

### Craniofacial dysmorphology in Dp1Tyb mice with intact Eustachian tube morphology

We observed that Dp1Tyb mice have craniofacial defects, consistent with recent reports of skeletal phenotyping in Dp1Tyb mice that identified abnormally shaped skulls^23,24^. As for other OM mouse models we analysed the craniofacial defects of the Dp1Tyb mice. 8- week-old Dp1Tyb mice appeared to have an abnormally shaped head with short neck, low set ears and rounded pinna (Fig. 2e). To investigate the craniofacial phenotype further we compared measurements of the skulls of Dp1Tyb and wild-type littermates. We measured the length of the nasal and frontal bone, and the frontal bone width. We normalised the data to the full skull length and detected a significant difference only in the frontal bone width of Dp1Tyb mice compared to wild-type mice (p = 0.0016) (Fig. 2f). We also investigated whether skull shape changes in Dp1Tyb mice affects Eustachian tube morphology which might impact the development of OME. However, measurements of the bony part of the tube did not detect any significant difference between the length or the width of the Eustachian tube (Fig. 2g) and thus it is unlikely that eustachian tube function is contributing to OME in the Dp1Tyb mice.

Finally, we compared weights of Dp1Tyb mice to wild-type littermates at the age of 3, 4 and 8 weeks. There was a significant difference between male mutants and wild-type mice at the first two time points but not at 8 weeks (mean weight of males at: 3 weeks wild-type 10.34 g, mutants 5.13 g, p <0.0001; 4 weeks wild-type 17.22 g, mutants 12.65 g, p = 0.0376; 8 weeks wild-type 26.46 g, mutants 23.33 g, p = 0.0901) (Extended Data Fig. 2). By the age of 8 weeks the surviving Dp1Tyb male mice had an average weight not significantly different from wild-type mice.

### Localisation of OME loci using the DpTyb mapping panel

We next utilised the nested series of genomic duplications across the Dp1Tyb region of Mmu16 to localise the locus or loci involved with susceptibility to OME. For each line we performed histological analysis of the middle ears to assess the incidence of OME. Analysis of Dp2Tyb, Dp3Tyb and Dp9Tyb mice revealed OME in Dp3Tyb and Dp9Tyb mice (Extended Data Fig. 3a and b, Extended Data Table 1). However, no OM was observed in Dp2Tyb mice. Out of fourteen Dp3Tyb carriers one did not have OME (7%), six had unilateral (43%) and seven bilateral OME (50%). The number of Dp3Tyb mice with OME was significantly different from wild-type mice (p = 0.0002) (Extended Data Fig. 3b, Extended Data Table 1). Out of twenty-six Dp9Tyb mice thirteen (50%) did not have middle ear inflammation, nine (34.6%) had unilateral and four (15.4%) bilateral OME (Extended Data Fig. 3b, Extended Data Table 1) significantly different from wild-type mice (p = 0.0339). The ABR threshold of the Dp3Tyb mice compared to wild-type mice was significantly raised by 20 dB SPL (p = 0.0072) when using the data from the mouse without OM in the calculation, or by 23.3 dB SPL (p = 0.0049) when Dp3Tyb mice with normal middle ears are excluded (Extended Data Fig. 3c). There was no significant difference in the threshold for the Dp9Tyb mice. The ABR threshold of the Dp9Tyb mice compared to wild-type mice was raised by 17.9 dB SPL (p = 0.0631) when using the data from the mouse without OM in the calculation, or by 11.7 dB SPL (p = 0.3137) when Dp9Tyb mice with normal middle ears are excluded (Extended Data Fig. 3c). Thus, the data demonstrates that there is a major locus with highly penetrant OME in the Dp3Tyb interval, and a minor locus in Dp9Tyb.

We then proceeded to further localise the major locus within the Dp3Tyb interval. Dp4Tyb, Dp5Tyb and Dp6Tyb mice have duplications of three shorter regions that together cover the entire region duplicated in Dp3Tyb mice (Fig. 3a). OME was not observed in Dp4Tyb and Dp6Tyb mice (Fig. 3b, Extended Data Table 1). However, all Dp5Tyb mice had OME, 33% of them unilateral and 67% bilateral OME (Fig. 3b, Extended Data Table 1), significantly different from wild-type mice (p < 0.0001). In addition, we also investigated the OME phenotype of Ts1Rhr mice which have a duplication that is shorter than that in the Dp3Tyb mice by 8 genes, but encompasses all of the region duplicated in Dp5Tyb mice (Fig. 3a). All the Ts1Rhr mutants had OME, 33% of them unilateral and 67% bilateral OME (Fig. 3a and b, Extended Data Table 1), significantly different to wild-type mice (p = 0.0004). We tested both Dp5Tyb and Ts1Rhr mice for hearing loss by performing click-evoked auditory ABR. At two months of age, the Dp5Tyb carriers had mean thresholds elevated by 28.5 dB SPL (p = 0.0081) and Ts1Rhr mice elevated by 19 dB SPL (p = 0.0248) compared to wild-type mice (Fig. 3c).

**Figure 3.**
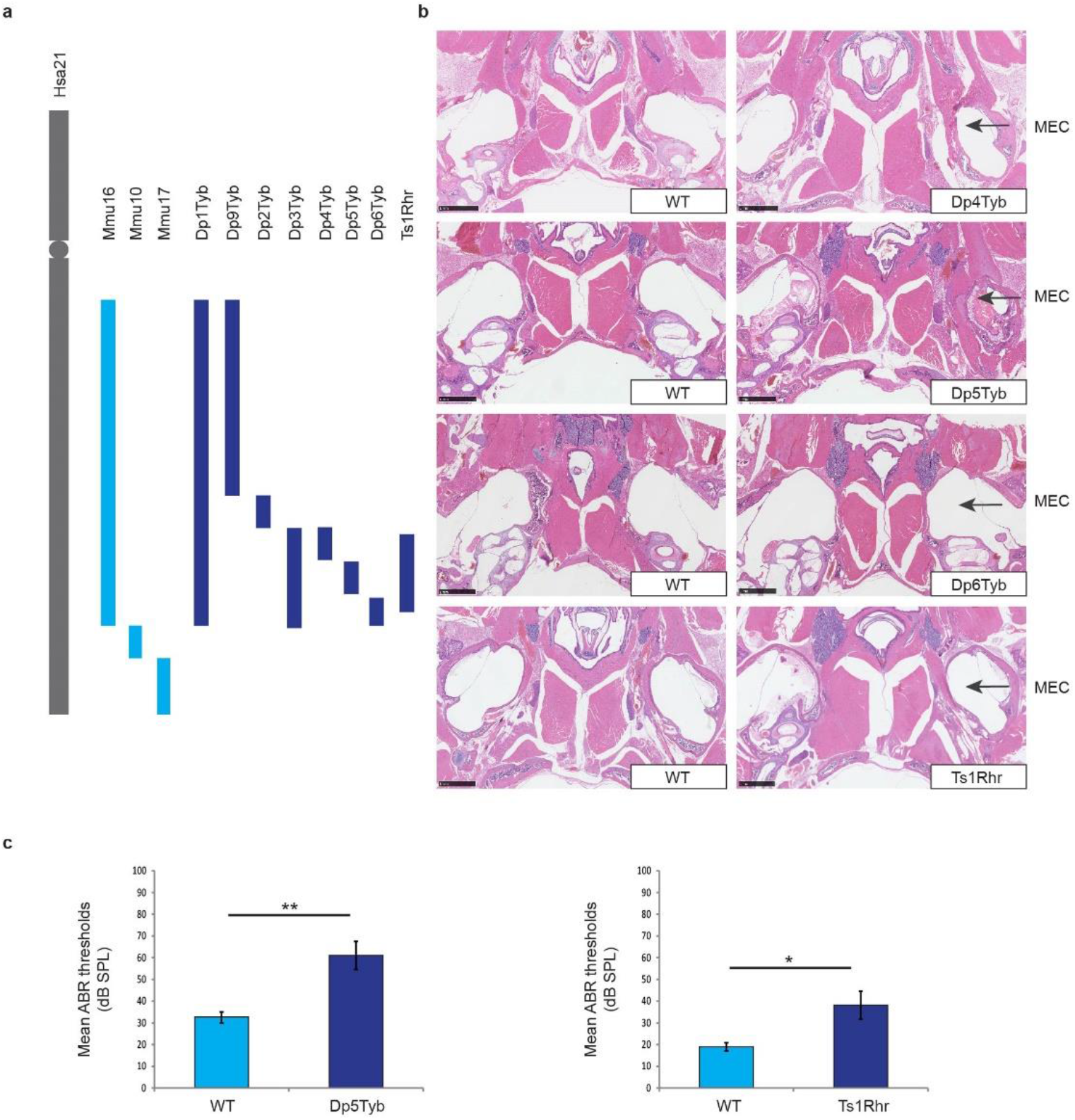
OM phenotypes of Dp4Tyb, Dp5Tyb, Dp6Tyb and Ts1Rhr mice. **a**, Representation of Hsa21 (in grey), the orthologous regions on Mmu10, Mmu17 and Mmu16 (in light blue) and the region of the duplication in Dp1Tyb, Dp9Tyb, Dp2Tyb, Dp3Tyb, Dp4Tyb, Dp5Tyb, Dp6Tyb and Ts1Rhr mice (in dark blue). **b**, Haematoxylin- eosin stained transverse sections through the middle ear of 2-month-old wild type (WT) and mutant mice showing middle ear inflammation in Dp5Tyb and Ts1Rhr mice. Note that the OM phenotype observed in the Ts1Rhr mouse displayed is unilateral in the left ear. MEC: middle ear cavity. Scale bars: 1 mm. **c**, Broadband click stimuli ABR thresholds in the right ears of 2-month-old wild-type (WT) and mutant (Dp5Tyb and Ts1Rhr) mice. The graph shows elevated mean thresholds in Dp5Tyb mice (n = 5 mice, 3 female and 2 male) compared to wild-type (WT) mice (n = 4 mice, 2 female and 2 male) and in Ts1Rhr mice (n = 5 mice, 2 female and 3 male) compared to wild-type (WT) mice (n = 5 mice, 1 female and 4 male). Bars: standard error of mean. P-values were determined using two-tailed t-test. * p≤0.05; ** p<0.01.

We carried out histological analysis on Dp7Tyb and Dp8Tyb mice that have duplications that between them cover the region duplicated in Dp2Tyb mice (Extended Data Fig. 4a) but neither, as expected, showed the presence of OME (Extended Data Fig. 4b, Extended Data Table 1). Finally, we assessed the middle ears of Dp12Tyb mice, that have a duplication of part of the region duplicated in Dp9Tyb mice, containing about half of the genes from the Dp9Tyb region (Extended Data Fig. 5a). None of the seven Dp12Tyb mice showed histological evidence of OME (Extended Data Fig. 5b, Extended Data Table 1). In conclusion, we have localised a major DS locus for OME within the Dp5Tyb region of Mmu16 (Fig. 4a and b). This region contains only 12 genes.

**Figure 4.**
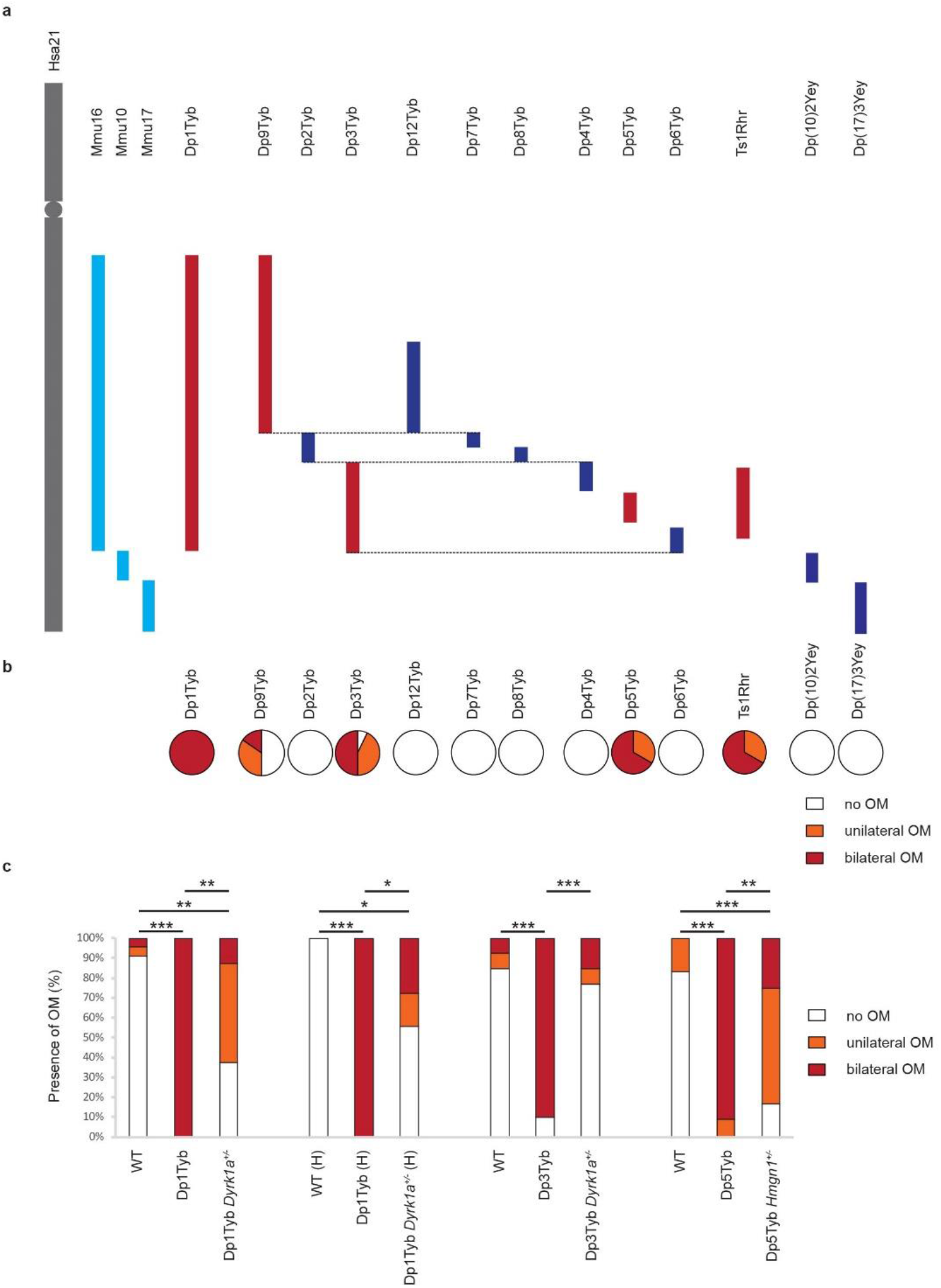
Localisation and verification of the major gene, *Dyrk1a*, involved in OME in DS. **a**, Representation of Hsa21 (in grey), the orthologous regions on Mmu10, Mmu17 and Mmu16 (in light blue) and the region of the duplication in all DpTyb mutants (in dark blue). **b**, The pie charts represent percentage of mutant mice with no effusion (in yellow) and with effusion in one (in orange) or in both ears (in red) for all the DpTyb lines tested along with Ts1Rhr, Dp2Yey and Dp3Yey. **c**, Normalisation of *Dyrk1a* dosage in Dp3Tyb mice leads to restoration of the wild-type phenotype. The graphs show the incidence of OM in wild-type (WT), DpTyb lines, and double mutant mice, carrying a DpTyb allele and heterozygous for a KO allele at either the *Dykr1a* or *Hmgn1* gene. First panel, mice produced from Dp1Tyb mice crossed to *Dyrk1a* knockout mice at the Francis Crick Institute - wild-type (WT), Dp1Tyb mice, and double mutant mice Dp1Tyb *Dyrk1a^+/-^*. Second panel, mice produced from Dp1Tyb mice crossed to *Dyrk1a* knockout mice at the MRC Harwell Institute (H) - wild-type (WT), Dp1Tyb mice, and double mutant mice Dp1Tyb *Dyrk1a^+/-^*. Third panel, mice produced from the Dp3Tyb mice crossed to *Dyrk1a* knockout mice in the Francis Crick Institute - wild-type (WT), Dp3Tyb mice, and double mutant mice Dp3Tyb *Dyrk1a^+/-^*. Fourth panel, mice produced from the Dp5Tyb mice crossed to *Hmgn1* knockout mice at the Francis Crick Institute - wild-type (WT), Dp5Tyb mice, and double mutant mice Dp5Tyb *Hmgn1^+/-^*. P-values were determined using two-tailed t-test. * p≤0.05; ** p<0.01; *** p<0.001.

### Normalisation of *Dyrk1a* dosage in Dp3Tyb mice leads to restoration of the wild- type phenotype

The 12 genes in the Dp5Tyb region represent candidates for the major OME gene in DS. Given its pleiotropic role in a number of cellular pathways, some of which might impact on epithelial inflammatory processes, we investigated if *Dyrk1a* dosage plays a key role in the development of otitis media in Down syndrome mouse models. We crossed Dp1Tyb mice to *Dyrk1a* knockout mice establishing replicate matings in two of the facilities we used to breed mice for this study – Francis Crick Institute and MRC Harwell Institute. Out of sixteen double mutants (Dp1Tyb *Dyrk1a*^+/-^) produced in the Francis Crick Institute six (37%) did not have OM, eight (50%) had unilateral and two (13%) bilateral OM which was significantly different from Dp1Tyb littermates (p = 0.0021). The wild-type phenotype was not completely restored as double mutants were significantly different from wild-type mice (p = 0.0022). Out of eighteen double mutants (Dp1Tyb *Dyrk1a*^+/-^) produced in the Harwell Institute ten had no OM (55%), three unilateral (17%) and five bilateral OM (28%) significantly different from Dp1Tyb littermate mice (p = 0.0110). Similar to the mice produced in the Francis Crick Institute there was a significant difference between the wild-type and double mutant mice (p = 0.0148) (Fig. 4c). We also carried out middle ear histology on *Dyrk1a* heterozygote knockout mice used for this study (four from the Francis Crick Institute and four from the MRC Harwell Institute) and found no evidence of OME. These findings suggest that *Dyrk1a* is one of the major genes contributing to the OM phenotype of the Dp1Tyb mice.

We next crossed *Dyrk1a* knockout mice to Dp3Tyb mice. Out of thirteen double mutants (Dp3Tyb *Dyrk1a*^+/-^) ten (77%) did not have OM, one (8%) had unilateral OM and two (15%) bilateral. Significantly fewer double mutant ears had OM compared to Dp3Tyb mice (p = 0.0001) and furthermore there was no significant difference in the phenotype between double mutants and wild-type mice (p = 0.5743) (Fig. 4c). Thus, normalising the gene dosage of *Dyrk1a* in Dp3Tyb mice showed a restoration of the wild-type phenotype demonstrating the role of *Dyrk1a* as a major gene causing the OM phenotype in Down syndrome.

In addition, we studied the candidature of the *Hmgn1* locus, which might also be expected to have pleiotropic roles impacting a number of organ systems. In this case we crossed Dp5Tyb mice to *Hmgn1* knockout mice and tested double mutants (Dp5Tyb *Hmgn1*^+/-^) carrying the duplication and heterozygous for the knockout allele. Out of twelve double mutants two (17%) did not have OM, seven (58%) had unilateral and three (25%) bilateral OM. We carried out histological analysis of the middle ears of the *Hmgn1* heterozygote knockout mice used for this study and found no OME. Despite a significant improvement in the frequency of OME between double mutants and Dp5Tyb mice (p = 0.0012), there was still a highly significant difference between double mutants and wild-type littermates (p = 0.0005) (Fig. 4c). Thus, the wild-type phenotype was not restored in the double mutants and we conclude that *Hmgn1*, unlike *Dyrk1a*, is not the major gene contributing to the OM phenotype of the Dp5Tyb mice.

### Expression of DYRK1A and associated pathway members of DYRK1A in middle ear epithelial cells, fluid and blood

Following the identification of the *Dyrk1a* gene as a major locus involved in Down syndrome OME we proceeded to investigate by immunohistochemistry the expression of the protein and associated pathway members in the middle ear in order to better understand the pathological mechanisms at the middle ear epithelium. DYRK1A was localised in the middle ear epithelial cells in both wild-type and mutant tissues. We detected a one-third increase, as expected, of the number of cells positive for DYRK1A in Dp3Tyb mice (56.6%) compared to wild-type littermates (36.4%) and a similar increase in Dp5Tyb mice (47.2%) compared to 28.6% in wild-type littermates (Fig. 5a and b). Given the potential cross-talk of DYRK1A signaling to the TGFβ pathway^25,26^ and the known involvement of TGFβ signalling in OME^8,9^, we studied the expression of pSMAD2 and SMAD3, members of the TGFβ family, in middle ear epithelial cells. We detected a significant increase in the number of cells positive for pSMAD2 (p = 0.0001, p = 0.0011) and SMAD3 (p = 0.0007, p = 0.0093) in Dp3Tyb and Dp5Tyb mice compared to wild-type littermates. Importantly, the number of cells positive for pSMAD2 and SMAD3 was restored to wild-type levels in double mutants (p = 0.6171, p = 0.8413) (Fig. 5a and b). Regulatory T cells/T helper cells (Treg/Th17) imbalances are reported in OME patients and mouse models ^27^. TGFβ is required in combination with inflammatory cytokines, such as IL-6, for the initial differentiation of Th17 and Treg cells^28^. DYRK1A is also known to regulate the differentiation of these cells^29^. We therefore examined expression of IL-6 in middle ear epithelial cells and recorded a significant increase in the number of cells positive for IL-6 (p = 0.0013, 0.0041), but not IL-10, in Dp3Tyb and Dp5Tyb mice compared to wild-type littermates (p = 0.4427, p = 0.3687). In addition, we found no significant difference between the number of cells positive for IL-6 in double mutants compared to wild-type mice (p = 0.2160) (Fig 5a and b). Furthermore, we employed Meso Scale Discovery Immunoassays to analyse the levels of several of those cytokines in middle ear fluids and bloods. We detected significantly more IL-6 in ear fluid compared to both wild-type (p = 0.0019) and mutant (p = 0.0042) blood samples (Fig. 5d). Levels of IL-17 and IL-21 (products of Th17 cells) were higher in ear fluid compared to wild-type (p = 0.0043, p = 0.0143) and Dp5Tyb (p = 0.0120, p = 0.0142) blood. The levels of IL-10 (product of Treg cells) in ear fluid, on the other hand were not significantly different from both blood samples (p = 0.7497, p = 0.1244) (Fig. 5d) and this was confirmed by immunohistochemistry analysis.

**Figure 5.**
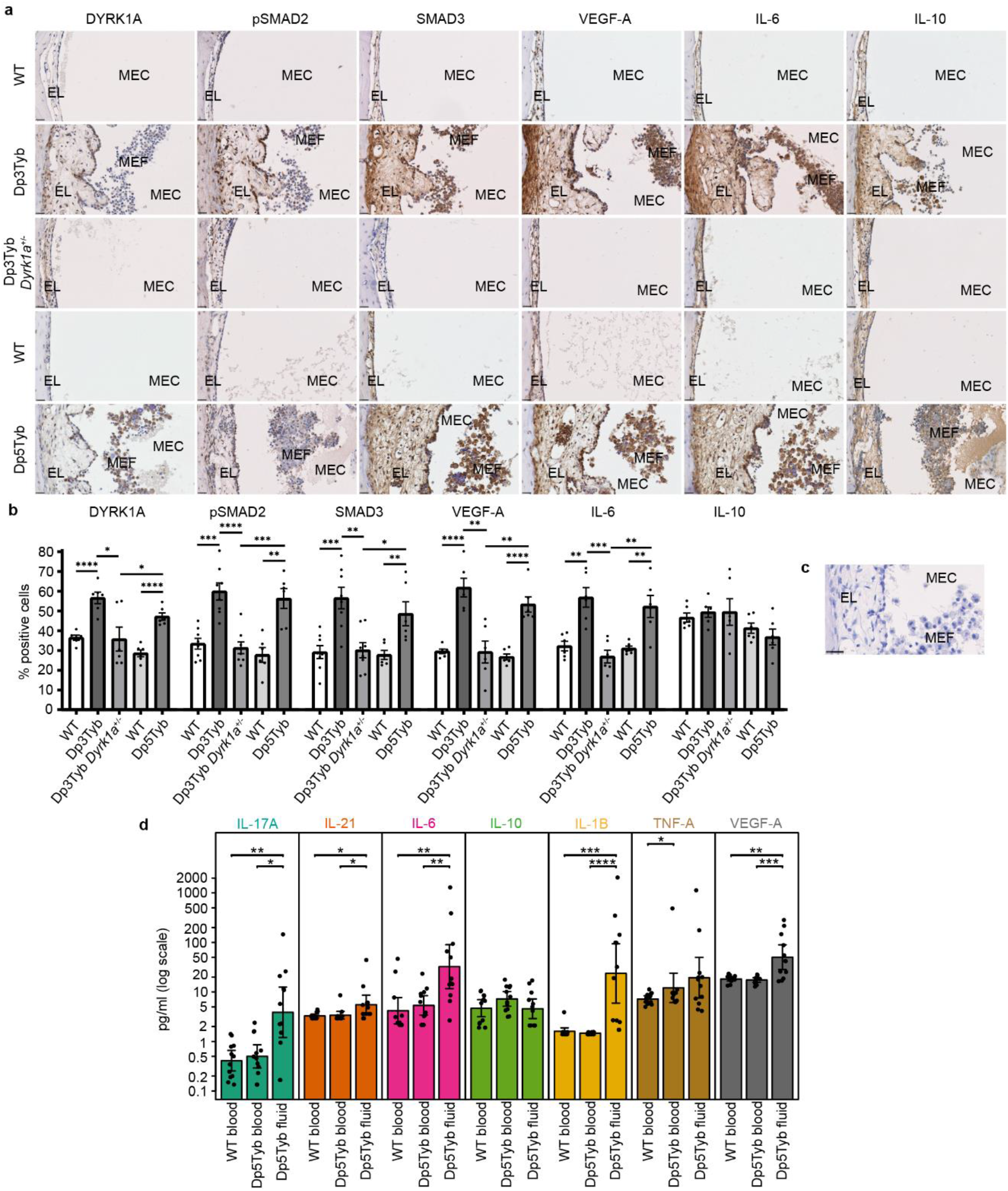
Expression of DYRK1A and associated pathway members. **a**, Immunohistochemical staining of transverse sections through the middle ear of mutant and wild-type (WT) mice with antibodies against DYRK1A, phospho SMAD2 (pSMAD2), SMAD3, VEGF-A, IL-6 and IL-10. Scale bar = 25 µm. EL: epithelial lining; MEC: middle ear cavity; MEF: middle ear fluid. **b**, Bar graphs of the immunohistochemistry shown in a, represented as percentage of middle ear epithelial cells that express the protein. To quantify the results middle ear epithelial cells were counted in six ears from each genotype for each antibody except for pSMAD2 and SMAD3 for the Dp3Tyb, Dp3Tyb *Dyrk1a^+/-^* mice and wild-type littermates (WT) where the cells were counted in eight ears. Error bars show standard error of the mean (SEM). Significance was established using two-tailed unpaired t-tests. P-values were denoted according to **** < 0.0001, *** < 0.001, ** < 0.01, * < 0.05. **c**, Negative control (no primary antibody) for the immunohistochemistry shown in (a). **d**, Mesoscale Discovery (MSD) assay using blood from Dp5Tyb mice (n = 12 mice, 11 female and 1 male), blood from wild-type (WT) littermates (n = 12 mice, 8 female and 4 male) and middle ear fluid from Dp5Tyb mutants (n = 12 mice, 11 female and 1 male). The results presented in the graphs are from two independent experiments for all antibodies. Each point corresponds to a single sample which has been measured repeatedly and averaged (see data pre- processing in methods). For each group in each panel the height of the coloured bar indicates the mean and whiskers extend to +/- 2 SEM. For each pairwise comparison in a panel a two-sided Mann–Whitney–Wilcoxon (MWW) test was performed and raw p- values were denoted according to: **** < 0.0001, *** < 0.001, ** < 0.01, * < 0.05. To control for multiple testing, we applied the Benjamin-Hochberg procedure to the complete set of 24 p-values arising from all the MWW tests across all assays and pairs of groups. Rejecting the null in all starred cases (*, **, *** and ***) controlled the FDR below 5%.

We also studied the involvement of DYRK1A in the VEGF pathway as VEGF levels in the middle ear are raised in OME, contributing to vascular leak and effusion into the middle ear cavity^4^. There were notably more cells positive for VEGFA in the middle ear epithelial cells in Dp3Tyb and Dp5Tyb mice compared to wild-type littermates (p < 0.0001, p < 0.0001). Moreover, the expression of VEGF was restored to wild-type levels in double mutants (p = 0.9545) (Fig 5a and b).

### Upregulation of hypoxia and inflammatory genes in Dp1Tyb and Dp5Tyb mice

The existence of hypoxia and elevated levels of *Hif1*α and HIF responsive genes *Vegfa*, *Il-1β* and *Tnfα* in middle ear fluids have been previously reported for a number of mouse models of OM: *Jeff*, *Junbo*^4^, and *edison* mice^5^. We therefore studied expression levels of these genes in blood and ear fluid from Dp1Tyb and Dp5Tyb mice. We first examined expression by RT-qPCR in blood samples from wild-type (n = 25 mice, 13 female and 12 male) and Dp1Tyb (n = 16 mice, 9 female and 7 male) mice and ear fluid from Dp1Tyb mutants (n = 17 mice, 10 female and 7 male). Relative to a baseline control of wild-type white blood cells, the inflammatory cells that accumulate within the middle ear fluid of Dp1Tyb mice showed elevated expression of *Hif1a* (30-fold; p <0.0001), *Vegfa* (260- fold; p <0.0001) and *Tnfα* (16-fold; p <0.0001) but not *Il-1β* (2-fold; p = 1) (Extended Fig. 6 and Extended Data Table 2). There was no significant difference in the expression levels between wild-type and mutant white blood cells (Extended Data Table 2). We then carried out similar analysis using wild-type (n = 13 mice, 8 female and 5 male) and Dp5Tyb (n = 19 mice, 9 female and 10 male) mice and ear fluid from Dp5Tyb mice (n = 9 mice, 4 female and 5 male). We recorded very similar results with elevated expression of *Hif1a* (11-fold; p < 0.0001), *Vegfa* (251-fold; p <0.0001) and *Tnfα* (59-fold; p <0.0001) but not *Il-1β* (2-fold; p = 0.8830) in ear fluid compared to wild-type blood samples (Extended Fig. 6 and Extended Data Table 2). In addition, using Meso Scale Discovery Immunoassay we found significant increases in protein levels of VEGF-A (p = 0.0056, p = 0.0009) and IL-1β (p = 0.0001, p < 0.0001) in ear fluid from Dp5Tyb mice compared to wild-type and Dp5Tyb blood samples (Fig. 5d). The increases in protein levels of VEGF-A are consistent with the results from immunohistochemistry described above.

**Figure 6.**
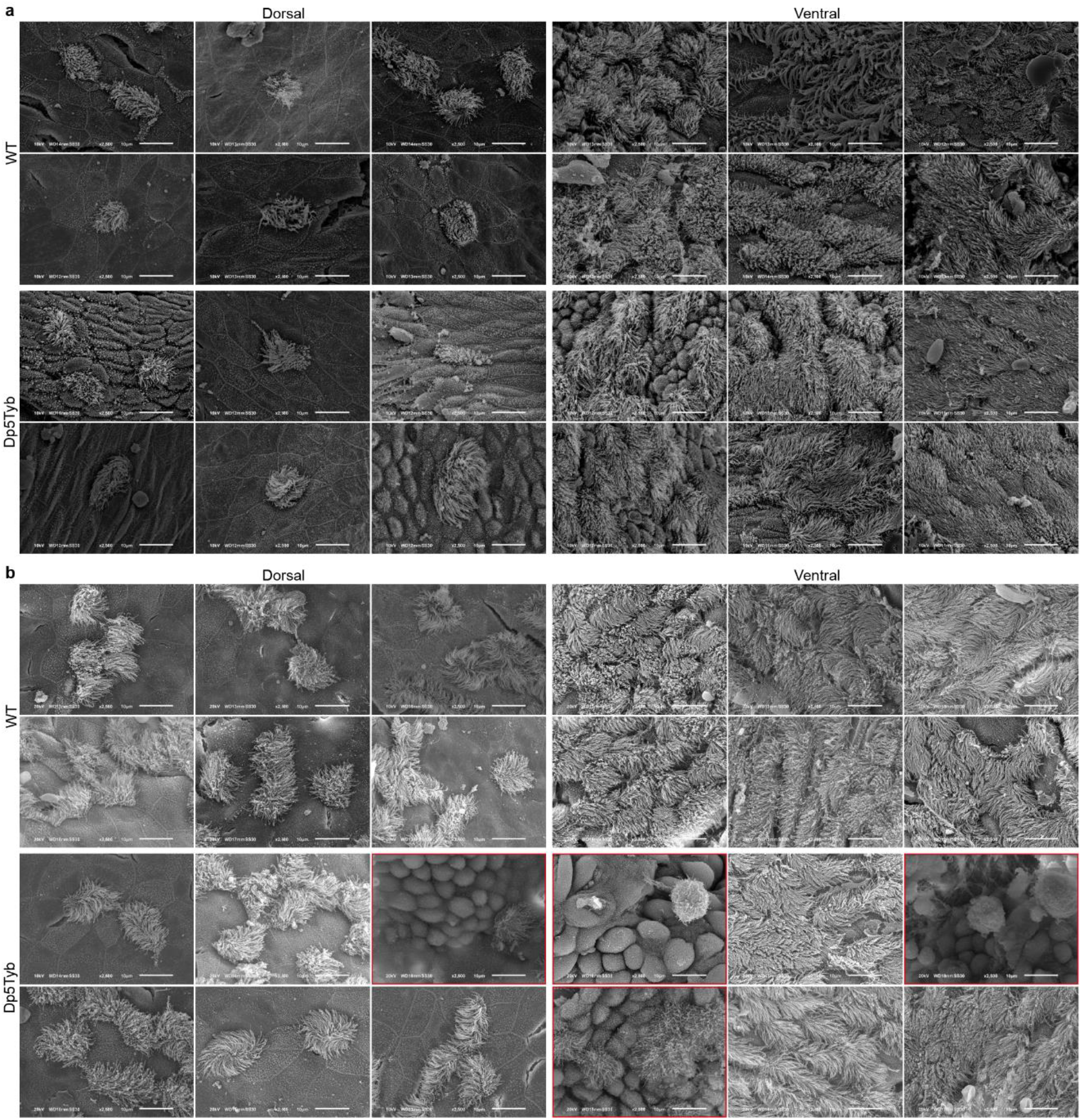
Cilia loss in the middle ear cavities of Dp5Tyb mice and littermate controls. Scanning electron microscopy (SEM) images from the dorsal and ventral region of the middle ear of **a**, 2-week-old and **b**, 2-month-old Dp5Tyb mice and wild-type (WT) littermates. Images were taken of both ears of each mouse (2-week-old Dp5Tyb n = 3 mice, 2 female and 1 male, WT n = 3 mice, 2 female and 1 male; 2-month-old Dp5Tyb n = 6 mice, 3 female and 3 male, WT n = 4 mice, 2 female and 2 male). Representative SEM images of middle ears were chosen for the panels. For the 2-week-old time point the dorsal and ventral images of wild-type ears and the dorsal images of the Dp5Tyb ears are from three separate mice. The 2-week-old ventral images of the Dp5Tyb ears are from two separate mice. For the 2-month-old time point the dorsal images of wild-type ears are from six different ears of five mice, the ventral images of wild-type mice - six different ears from three mice. Both, dorsal and ventral images of middle ear images of 2-month-old Dp5Tyb mice represented in the figure, at the time when the mice have fully developed OM, are from six different ears from five mice (for the dorsal) and from six different ears for four mice (for the ventral). Loss of cilia in the middle ear of epithelial cells was detected only in the ears of 2-month-old mutants (highlighted in the figure with red borders) but not in 2-week-old Dp5Tyb mice. A complete lawn of cilia is no longer present in the highlighted ventral images of the mutant mice. Images taken at x2500 magnification. Scale bar = 10 µm.

### Cilia loss in the middle ear epithelium of Dp3Tyb and Dp5Tyb mice

The middle ear epithelium arises during development from two distinct lineages^30^. Dorsally in the middle ear cavity the epithelium is derived from neural crest, whereas ventrally and around the Eustachian tube the epithelium is endodermal in origin. The ventral epithelium develops cilia, which play a role in middle ear clearance, whereas in the dorsal region few cilia are present. DYRK1A has recently been implicated in ciliogenesis^31^, so we sought to uncover the effects of *Dyrk1a* triplication on ciliogenesis in the middle ear and the role that this pathway might play in OME. We used SEM to study cilia coverage in wild-type and Dp5Tyb mice in ventral and dorsal regions of the middle ear at 2 weeks of age, before OME develops, and also at 2 months. At 2 weeks, in Dp5Tyb mice, compared to wild-type, cilia development was normal with the expected distribution of cilia in both ventral and dorsal regions (Fig. 6a). Similarly, in Dp3Tyb mice, cilia development was normal apart from one mouse which showed some evidence of cilia loss in the dorsal region (Extended Fig. 7a). In contrast, at 2 months of age in Dp5Tyb mice, in comparison with wild-type, we saw extensive cilia loss in the ventral region in one mouse, and extensive cilia loss in the dorsal region in 3 mice (Fig. 6b). A similar picture was found in Dp3Tyb mice at 2 months of age, where in the ventral region 2 mice showed extensive cilia loss, and in the dorsal region 4 mice had lost the uniform carpet of ciliated cells that characterizes the region of the middle ear (Extended Fig. 7b). The normal cilia development in both Dp5Tyb and Dp3Tyb mice at 2 weeks of age indicates that there is no direct effect of three copies of *Dyrk1a* (or any other gene in these regions) on cilia development. The extensive cilia loss observed in 2-month old Dp5Tyb and Dp3Tyb mice is thus likely secondary to the inflammatory OME that develops and is not a direct result of trisomic effects on cilia development. Nevertheless, the cilia loss that is observed may be a contributory factor to the development of OME in DS.

### Expression of duplicated genes from the Dp5Tyb region in children with Down syndrome

Finally, we studied the twelve genes from the Dp5Tyb region to test if their expression is increased in samples from children with DS compared to unaffected parents. qPCR results comparing saliva DNA from six paired Down syndrome children with OM and unaffected mothers revealed that out of the twelve genes tested the gene with the highest fold difference, 1.54, was *DYRK1A*. There was a significant difference in the expression of *DYRK1A* in saliva samples from children compared to their mothers (p = 0.004) (Extended Data Table 3).

## Discussion

Otitis media with effusion is a very common middle ear disease in non-syndromic children^2^ but is even more common in children with Down syndrome^1^. Whereas there has been significant progress using mouse models in identifying genes and genetic pathways involved in COME and that might contribute to susceptibility to the disease in the human population^4–9^, we remain ignorant of the underlying genetic loci that are involved with otitis media in DS. The localisation and identification of the genetic loci for COME in DS will enable us to develop a molecular understanding of the pathological mechanisms. Moreover, we can relate the loci involved in DS to genes and pathways identified from mouse models of COME and potential loci that are involved in susceptibility in the human population^11^. Collectively, insights will emerge from these studies that may lead to improved treatments of this serious and common condition.

To identify critical regions and genes that contribute to genetic susceptibility to COME in DS we used a panel of mouse lines with complete and partial duplications of the gene regions from the three mouse chromosomes that are orthologous to Hsa21. We tested each of the mouse lines for a middle ear OM phenotype and for hearing loss. We found that the Dp1Tyb line demonstrates a fully penetrant bilateral OME phenotype with a highly cellular exudate arising at weaning age. We did not observe COME in either the Dp2Yey or Dp3Yey lines demonstrating that Hsa21-orthologous genes on Mmu10 and Mmu17 are not involved. However, we employed the nested set of duplications covering Dp1Tyb to localise the COME genetic loci to the Hsa21-orthologous region of Mmu16.

We identified a major highly penetrant locus predisposing to COME in Dp3Tyb, which was narrowed down by the examination of Dp4Tyb, Dp5Tyb and Dp6Tyb lines. Only Dp5Tyb mice showed COME, which is consistent with the observation of COME in the Ts1Rhr line which has a duplication of a region that encompasses and is slightly larger than that duplicated in Dp5Tyb mice. Analysis of Dp9Tyb mice suggests the presence of a minor locus on mouse chromosome 16. We suggest that the complete penetrance observed in the Dp1Tyb line may arise through additive effects of the major locus in the Dp5Tyb region and a minor locus within the DpTyp9 region. Overall, we conclude that a major, highly penetrant locus conferring susceptibility to COME lies within the Dp5Tyb region. This small region contains only 12 genes: *Dyrk1a, Kcnj6, Kcnj15, Erg, Ets2, Psmg1, Brwd1, Hmgn1, Wrb, Lca5l, Sh3bgr* and *B3galt5*^20^.

Various trisomic mouse models of DS have been examined for evidence of COME^13,18,19^. Our finding of a major locus in the Dp1Tyb region, and localised to the Dp5Tyb region, is consistent with the COME observed in the Ts65Dn^13^ and Dp(16)1Yey mouse strains^19^. Two studies have separately reported that they find no evidence of COME in Tc1 transchromosomic mice that carry an almost complete copy of human chromosome 21^18,19^. The human 21 Tc1 chromosome however, carries two deletions and it has been surmised that genes carried within these two regions may contribute to COME susceptibility. One of these regions contains five genes, *CXADR, BTG3, C21orf91, CHODL* and *TMPRSS15* and lies within the Dp9Tyb region. The other region is contained within Dp2Tyb and encompasses *MIS18A, MRAP, URB1, EVA1C, C21orf59, SYNJ1, PAXBP1, C21orf62, OLIG2, OLIG1, IFNGR2, AP00295.1, IL10RB, IFNAR1, IFNGR2, TMEM50B, DNAJC28, GART, SON, DONSON, ATP5PO, CRYZL1, ITSN1, MRPS6, SLC5A3, KCNE2, SMIM11A, FAM243A, KCNE1, RCAN1, CLIC6* and *RUNX1*.

Thus, these two deleted regions do not overlap with the Dp5Tyb region, suggesting that other factors are impacting on the development of COME in the Tc1 mouse. These factors could include genetic background and environmental effects, and a critical aspect of the current study is that all the DpTyb mouse lines are maintained on the same C57BL/6J genetic background. Moreover, our findings on the presence of COME in the different Dp lines are reproducible across two centres (Francis Crick Institute and Harwell Institute).

To identify the gene or genes within the Dp5Tyb region which are responsible for OME we crossed relevant Dp lines to gene knockouts in order to normalize gene copy number and to assess whether the wild-type phenotype was restored in progeny carrying two gene copies. Crosses of Dp1Tyb and Dp3Tyb to *Dyrk1a* knockouts demonstrated that restoring the normal copy number led to a significant reduction in OME. It is noteworthy that in the Dp3Tyb cross we saw a dramatic improvement in the middle ear phenotype such that the ‘two gene’ progeny were not significantly different from wild-type. It is possible that the improvement in Dp1Tyb double mutants was less dramatic given the continued triplication of the minor Dp9 locus in the disomic progeny from the Dp1Tyb cross. We also tested the *Hmgn1* locus and data from the Dp5Tyb *Hmgn1* knockout cross suggested that although the double mutants produced from the cross have reduced incidence of OM, *Hmgn1* is not the main gene as the wild-type phenotype was not restored. Overall, the findings demonstrate that *Dyrk1a* is a major gene involved with the development of OME in DS.

In order to investigate the pathological mechanisms of *Dyrk1a* trisomy that lead to OME, we investigated the known pathways in which *Dyrk1a* is involved. This includes firstly its impact on Hedgehog (Hh) signalling^26^ and cross-talk with TGFβ signalling^25^ which is already known to be involved with OME^8,9^. We therefore assessed the levels of pSMAD2 and SMAD3 from the TGFβ pathway in middle ear epithelium and found that they were significantly raised in Dp3Tyb mice, but restored to normal wild-type levels in double mutants. TGFβ is a key contributing factor for the reciprocal differentiation of Th17 versus Treg cells and the fate of these cells depends on the presence of proinflammatory cytokines like IL-6. TGFβ in combination with IL-6, a proinflammatory cytokine, can induce Th17 differentiation from naïve CD4+ T cells and inhibit the differentiation of Treg cells. In the absence of proinflammatory cytokines TGFβ induces the differentiation of Treg cells^28^. More recently DYRK1A was also identified as part of a pathway that suppresses the development of Treg cells^29^. In Dp5Tyb and Dp3Tyb mice, we recorded high levels of pSMAD2 and SMAD3 (which operate downstream of the TGFβ ligands) and high levels of IL-6 in the middle ear epithelial cells. This potential cross-talk of DYRK1A signaling and TGFβ pathway in the middle ear could result in differentiation of naïve CD4+ T cells into Th17 cells. Cytokines expressed by Th17 cells are known to have a role in the pathogenesis of inflammatory disease^32^. One of those cytokines is IL- 17 which is reported to be overexpressed in inflamed lung endothelial cells^33^. For that reason, we assessed the levels of IL-17 and detected a significant increase of the protein in Dp5Tyb middle ear fluid compared to wild-type bloods.

Second, DYRK1A positively regulates VEGF-dependent NFAT transcription in endothelial cells and when silenced reduces intracellular Ca2+ influx in response to VEGF along with a reduction in VEGFR2 receptors^34^. Thus, DYRK1A overexpression may enhance the VEGF response and exacerbate vascular leak from endothelial cells in the middle ear epithelium that arise from raised levels of VEGF ^4^. Indeed, we show, as for other mouse mutants with OME, that levels of VEGF in the middle ear epithelium are significantly raised in Dp3Tyb mice and levels are restored to wild-type in double mutants. Moreover, we found that the expression levels of *Hif1a* are raised in middle ear exudate in Dp1Tyb mice. These findings indicate that the middle ears of the Dp1Tyb mice are hypoxic, potentially also contributing to upregulation of VEGF. These results highlight similarities between the downstream molecular pathology identified in the Dp1Tyb line and previously characterised mouse mutants with OME^4,5,9^. This hypoxic response has also been observed in human middle ear effusions by transcript analysis and the determination of VEGF protein titres that were found to be significantly raised^35^. It has been surmised that the hypoxic response and upregulated VEGF signalling found in mouse models and humans underlie the vascular leak, angiogenesis and lymphangiogenesis that are hallmarks of OM and identifies VEGF and VEGF receptors as potential targets for the treatment of OME^4^.

Third, it has recently been reported that DYRK1A plays a role in ciliogenesis^31^. We therefore explored the histopathology of cilia in the middle ear, which may play a role in clearance of middle ear effusion and thus contribute to the onset and severity of OME. In Dy5Tyb and Dp3Tyb mice at 2 months of age, in both the dorsal and ventral regions of the middle ear we found in that there was significant loss of ciliated epithelium. However, this was not observed at 2 weeks of age, indicating that likely the observed cilia loss at 2 months of age is secondary to the development of OME and not a direct result of *Dyrk1a* trisomy. We had previously observed loss of ciliated epithelium in the middle ears of the *Jeff* mutant mouse, which carries a mutation in the *Fbxo11* gene and displays chronic otitis media^36^. Other genes involved in cilia function and which are triplicated in DS have been studied for their role in DS phenotypes. Ts1Rhr mice were reported to have ventricular enlargement accompanied by ependymal cilia beating deficiency. The *Pcp4* gene (Purkinje cell protein 4) lies within the Ts1Rhr trisomic segment and regulates calmodulin and intracellular calcium concentration. Restoration of *Pcp4* to two copies rescued both phenotypes in Ts1Rhr mice^37^ suggesting a key role of *Pcp4* in cilia function. However, *Pcp4* is present in three copies in Dp6Tyb mice, which do not develop OM. Trisomy and increased levels of the Pericentrin gene (*PCNT*) have been associated with disruptions to centrosomal trafficking and cilia development^38^. *Pcnt* is on Mmu10, in the Dp(10)2Yey segment and we found that Dp(10)2Yey mice do not develop middle ear inflammation ruling out the involvement of Pericentrin in OME is DS. In summary, there are a variety of pathophysiolocial impacts of *Dyrk1a* in three copies and the evidence suggests that a number of pleiotropic phenomena are contributing to the development of OME in DS.

Three copies of *Dyrk1a* has been implicated for some of the phenotypes found in Down syndrome such as neurogenesis and brain development defects, neurodegeneration and congenital heart defects^39–41^. It has been reported that normalising the gene dosage of *Dyrk1a* in the Ts65Dn Down syndrome mice rescues some of the Alzheimer’s disease phenotypes^42^. Mice with *Dyrk1a* over expression were found to develop heart failure^43^. Moreover, normalizing the gene dosage of *Dyrk1a* from 3 to 2 copies in Dp1Tyb mice reverses the congenital heart defects observed in this strain^41^. We now show that *Dyrk1a* is involved in another major disease feature of DS, and it is intriguing to speculate that many phenotypes of DS emerge from a few critical genes on human chromosome 21, some of which have widespread pleiotropic impact on a number of organ systems. Indeed, this is supported by a report of a 5.5 year old boy with features of DS, including craniofacial dysmorphism and development delay, found to be carrying a microduplication that encompassed only the DYRK1A gene. Whether or not the boy was affected by OME at that age was not reported^44^.

A number of inhibitors of DYRK1A have been recently identified and used to study the molecular mechanisms of DYRK1A action. Harmine was used to identify DYRK1A as a regulator of Treg cell differentiation^29^. The small-molecule Wnt pathway inhibitor lorecivivint (SM04690) was reported to modify the Wnt pathway through the inhibition of CLK2 and DYRK1A^45^. A recent study examining the therapeutic potential of another small-molecule Wnt pathway inhibitor, SM04755, for the treatment of tendinopathy identified DYRK1A as a target for the inhibitor^46^. Most recently a natural product from herbal plants, aristolactam BIII, was also identified as a novel DYRK1A inhibitor and proposed as a potential therapy for Down syndrome-related disorders^47^. Moreover, Leucettinibs, a new class of DYRK/CLK kinase inhibitors, have recently been reported^48^ and leucettinib-21 appeared to cause a partial reversal of CHD in Dp1Tyb mouse embryos^41^. The middle ear is potentially directly accessible for topical treatment^49,50^ and the identification of *Dyrk1a* as a major gene contributing to OME in DS provides a novel route for the treatment of this condition. Suppressing the activity of DYRK1A by localized delivery of inhibitors to the middle ear cavity in Down syndrome patients should be explored as a potential strategy for future treatment of OME.

## Materials and Methods

### Mice

Mice carrying the Dp(16Lipi-Zbtb21)1TybEmcf (Dp1Tyb), Dp(16Mis18a- Runx1)2TybEmcf (Dp2Tyb), Dp(16Mir802-Zbtb21)3TybEmcf (Dp3Tyb), Dp(16Mir802-Dscr3)4TybEmcf (Dp4Tyb), Dp(16Dyrk1a-B3galt5)5TybEmcf (Dp5Tyb), Dp(16Igsf5-Zbtb21)6TybEmcf (Dp6Tyb), Dp(16Lipi-Hunk)9TybEmcf (Dp9Tyb), Dp(10Prmt2-Pdxk)2Yey (Dp(10)2Yey), Dp(17Abcg1-Rrp1b)3Yey (Dp(17)3Yey), Dp(16Cbr1-Fam3b)1Rhr (Ts1Rhr) and *Dyrk1a*^tm1Mla^ (*Dyrk1a* KO) alleles have been described previously^20,21,51,52^. The generation of mice carrying Dp(16Mis18a- Il10rb)7TybEmcf (Dp7Tyb), Dp(16Ifnar1-Runx1)8TybEmcf, Dp(16Krtap24-1- Hunk)12TybEmcf (Dp12Tyb) and *Hmgn1*^em1Tyb^ alleles will be described elsewhere (Lana-Elola, Watson-Scales, Fisher and Tybulewicz manuscript in preparation). The *Hmgn1*^em1Tyb^ allele has a 360 bp CRISPR/Cas9-generated deletion (16:96127399 - 16:96127758) which removes all of exon 1, intron 1 and part of exon 2. They were bred and maintained at the Mary Lyon Centre, MRC Harwell and at the Francis Crick Institute and were housed in specific-pathogen free conditions. All mouse strains were maintained on a C57BL/6JNimr background. Genotyping was carried out using custom probes (Transnetyx). The humane care and use of mice in this study was under the authority of the appropriate UK Home Office Project License.

### Histology

Heads from 3-, 4-, 8- and 16-week old mice were collected and fixed in 10% buffered formaldehyde, decalcified and embedded in paraffin following routine procedures. 5 µm- thick sections were obtained and stained with haematoxylin and eosin for morphological observations.

### Immunohistochemistry

Paraffin sections of mouse heads from 8- and 16-week-old mutant and wild-type mice were de-waxed and rehydrated following routine procedures. Endogenous peroxidase was blocked with 3% hydrogen peroxide and sections were incubated overnight with primary antibodies against: DYRK1A, 1:100 (ab65220, Abcam); phospho SMAD2 (Ser465/467), 1:100 (AB3849-I, Merck Millipore); SMAD3, 1:100 (06-920, Merck Millipore); VEGFA, 1:200 (AB1876-I, Merck Millipore); IL-6, 1:100 (ab6672, Abcam); and IL-10, 1:100 (ab217941, Abcam). The Vectastain Elite ABC HRP kit (PK-6101, Vector Laboratories) kit was used according to the manufacturer’s instructions for all of the antibodies. For development of the signal, the DAB+ chromogen system was used (K3468, DAKO). Counterstaining was carried out with haematoxylin.

### Meso Scale Discovery Immunoassay

A plate was designed to contain analytes of interest from the U-PLEX Biomarker Group 1 (ms) Assays (catalogue number K15069L-1). U-PLEX Mouse analytes: IL-1β, IL-6, IL-10, IL-17A, IL-21, TNF-α, and VEGF-A. The MSD plate was coated and loaded following the manufacturer’s instructions. Blood samples from eleven female and one male 8-week-old Dp5Tyb mice and eight female and four male wild-type littermates were collected via retro-orbital bleed into brown z-gel tubes (no anti-coagulant). Serum was isolated by allowing the blood to clot at room temperature for 1 hour before centrifuging (16000 x g, 4°C, 5 min). The supernatant (serum) was loaded neat into the plate (25 µl/well). To obtain middle ear fluids from eleven female and one male Dp5Tyb mice the tympanic membrane was removed with forceps. The fluid was collected from both ears of each mouse, added to 50 µl ice cold PBS, vortexed for 30 seconds and centrifuged (500 x g, 4°C, 10 min) to pellet cells and debris. The supernatant was removed and added to the plate neat (25µl/well). The standard curve was created using 4-fold serial dilutions of the calibrators in Diluent 41, and diluent alone as the 8th standard.

### Auditory brainstem response (ABR)

8-week old mice, from each genotype and gender, were anaesthetised and placed on a heated mat in a sound attenuating chamber. Acoustic stimuli were delivered to the right ear and ABR responses were collected, amplified and averaged using the BioSig software. Broadband click stimuli were presented at 90 dB SPL and gradually decreased in steps of 5 dB until a threshold was visually determined by the lack of replicable response peaks. The test was analysed, as previously described^53^.

### Skull morphology

Skulls from 8-week old mice were dissected, macerated in 1% potassium hydroxide and stained with alcian blue for cartilage and alizarin red for bone, following routine histological procedures. The skulls were stored in glycerol until measurements were taken. ImageJ software was used to measure the skull length, nasal bone length, frontal bone length and skull width. Allometric comparisons were performed against skull length with at least three mice of each genotype and sex.

### Scanning electron microscopy

Inner ears, dissected from two female and one male 4-week-old Dp1Tyb mice and wild-type littermates, were fixed, washed and decalcified, as previously described^54^. The organ of Corti was exposed, the ears were dehydrated, critical-point dried, sputter coated with gold/platinum and then viewed on a JEOL 6010 LV scanning electron microscope.

Middle ears from: two female and one male 2-week-old Dp5Tyb mice and wild-type litter mates, two female and one male 2-week-old Dp3Tyb mice and one female and two male wild-type littermates, three female and three male 2-month-old Dp5Tyb mice and two female and two male wild-type littermates, three female and three male 2-month-old Dp3Tyb mice and one female and two male wild-type littermates were prepared for imaging using the same methods as described for the cochleae, except no decalcification or fine dissection was required as the cilia are already exposed, and samples were additionally stained using osmium tetroxide. Images were taken at 2500x magnification.

### Real-time quantitative PCR (RT-qPCR) analysis of blood and middle ear fluid

Middle ear fluid and venous blood were collected from 8-week-old mice. For this study ear fluid from 10 female and 7 male mice; blood from 9 female and 7 male Dp1Tyb mice; blood from 13 female and 12 male wild-type mice was used. RNA from the blood samples was extracted using the Mouse RiboPure blood RNA isolation kit (1951, Ambion) and RNA from the ear fluid samples was extracted using RNeasy Plus micro kit (74034, Qiagen). For the Dp5Tyb mice, RNA was extracted from venous blood collected from nine female and ten male Dp5Tyb mice and eight female and five male wild type mice using the Maxwell RSC simplyRNA blood kit (AS1380), and from four female and five male Dp5Tyb ear fluids using the Maxwell RSC simplyRNA cells kit (AS1390). All mice were 8-weeks-old. After the RNA extraction cDNA was synthesised, RT-qPCR was performed and the data was analysed, as previously described^5^.

### qPCR analysis of saliva samples

Saliva samples were collected from six children with Down syndrome who underwent surgery for otitis media (4 male, 2 female; average age 5.7 years, range 0.5-14.8 years; 4 with COME, two with recurrent acute otitis media), and their unaffected mothers using Oragene saliva kits (DNA Genotek, Ottawa, Ontario, Canada). DNA samples were isolated from saliva according to the manufacturer’s instructions. Primers were designed for the 12 genes within the mouse region of interest and control gene *ACTB* using the human reference sequence (hg19). Each of the 12 DNA samples were used for qPCR on a Bio-Rad machine (Hercules, CA, USA) in triplicate using the SYBR Green PCR Master Mix (Thermo Fisher Scientific). Fold change was determined based on ΔCT values.

## Data analysis

We used the Chi-squared test to compare the difference between the observed and the expected number of the Dp1Tyb mice. Two-tailed t-test was used for comparing mean ABR thresholds, mucoperiosteal thickness, skull, and Eustachian tube measurements. We used two-tailed Fisher’s exact test for comparing mutant and wild-type mice for prevalence of OME. A value of p less than 0.05 was considered significant. Tukey *post hoc* multiple comparisons analysis using ANOVA of the delta CT means against the genotype, gene expression probe and sample collection type were calculated to determine qPCR expression differences with a family-wise confidence level of 95%.

For MSD data preprocessing we replaced missing values with LOD / 2 for the specific assay (Lower LODs in pg/ml are IL-17A = 0.3, IL-21 = 6.5, IL-6 = 4.8, IL-10 = 3.8, IL-1B = 3.1, TNF-A = 1.3, VEGF-A = 0.77) and applied logarithmic (base 2) transformation. We regressed out plate effects (subtract plate means and add global mean) and calculated the mean of data for each sample (12 samples have four technical replicates averaged from across two plates, and 24 samples have two technical replicates averaged from a single plate).

For each pairwise comparison in a panel a two-sided Mann–Whitney–Wilcoxon (MWW) test was performed and raw p-values were denoted according to (**** < 0.0001, *** < 0.001, ** < 0.01, * < 0.05). To control for multiple testing, we applied the Benjamin-Hochberg procedure to the complete set of 24 p-values arising from all the MWW tests across all assays and pairs of groups. Rejecting the null in all starred cases (*, **, *** and ***) controlled the FDR below 5%.

## Acknowledgments

The authors would like to thank the staff of the Mary Lyon Centre for animal husbandry, in particular Simona Oliveri, Elizabeth Joynson, Simon Gillard, Michelle Stewart and Heather Cater; Caroline Barker and Adele Austin for histology services; the genotyping core; Sara Johnson and Zsombor Szoke-Kovacs for the microCT scans; Lee Moir for the assistance using MSD plates and George Nicholson for the help with the statistical analysis of the data from the MSD plates. We also thank T Bootpetch for performing qPCR, and N Friedman, K Chan, M Scholes, S Haville and K Tholen for providing saliva DNA samples for study.

This work was supported by the Medical Research Council, UK (MC_U142684175 to S.D.M.B.), Wellcome Trust (grant numbers 080174, 098327 and 098328, to E.M.C.F. and V.L.J.T.), National Institutes of Health – National Institute on Deafness and Communication Disorders (R01 DC015004 to R.L.P.S.C.) and the Ponzio Research Accelerator Award from the Center for Children’s Surgery, Children’s Hospital Colorado (to B.W.H.). V.L.J.T. was supported by the Francis Crick Institute which receives its core funding from Cancer Research UK (FC001194), the UK Medical Research Council (FC001194), and the Wellcome Trust (FC001194). This research was funded in whole, or in part, by the Wellcome Trust. For the purpose of Open Access, the authors have applied a CC-BY public copyright licence to any Author Accepted Manuscript version arising from this submission.

## Author contributions

H.T., S.D.M.B., A.S., E.L-E, V.L.J.T and E.M.C.F. conceived and designed the experiments and interpreted the results. E.L-E., S.W-S., B.W.H., R.S-C. and D. G. were involved in the provision of resources. H.T., A.S., P.V., D.W., T.P., P.M., and A.P. performed the experiments, A.S., P.V., H.V.L., R.S-C. and D.N. contributed to analyzing the data, S.W. coordinated the management of the mouse colonies, H.T. and S.D.M.B. wrote the manuscript., A.S., E.L-E, V.L.J.T and E.M.C.F. were involved in correcting and editing of the manuscript.

